# Tracking multi-site somatic voltage dynamics via high-speed fiber photometry

**DOI:** 10.64898/2026.06.02.729189

**Authors:** S. Chakraborty, M. van Veghel, A. Tzanou, Z. Li, D. Torbin, E. Lowet

## Abstract

Investigating neural circuit dynamics across distributed brain regions in awake, behaving animals is crucial for understanding complex behavior. Genetically encoded voltage indicators (GEVIs) offer a powerful approach to tracking transmembrane voltage with high temporal and cellular specificity. However, scaling high-sensitivity GEVI recordings across multiple brain regions and multiple animals simultaneously remains a major technical challenge. Furthermore, it is unclear whether soma-targeted GEVIs - typically used for single-cell resolution imaging - can be effectively adapted for fiber photometry. Here, we show that a sCMOS-based widefield imaging system achieves sensitive dual-color multi-site fiber photometry using soma-targeted GEVI indicators with high temporal resolution. We validated this approach in the mouse hippocampal CA1, capturing theta (3-10Hz) and gamma (30-80Hz) rhythms and theta-gamma cross-frequency coupling. Additionally, we recorded high-frequency neural entrainment (>100 Hz) and somatic depolarization induced by electrical stimulation in CA1. Lastly, we tracked synchronized neural activity between the bilateral CA1s as well as multi-site dual-color imaging across CA1 and cortex simultaneously in three freely running mice. This work provides a scalable, accessible platform for high-speed optical electrophysiology in distributed neural circuits.

**Key points:**

- Implementation of a sCMOS-based widefield imaging setup for sensitive and scalable fiber photometry and cellular imaging.
- Demonstration of population multi-site and inter-animal voltage imaging with soma-targeted genetically encoded voltage indicators.
- Tracking of high-frequency gamma and high-gamma (>100Hz) neural entrainment

## Introduction

Monitoring neural population activity is essential for decoding brain function in both clinical settings and basic research. Traditionally, this has relied on extracellular recordings such as electroencephalography (EEG), local field potentials (LFP), and magnetoencephalography (MEG). While these methods provide a window into large-scale network dynamics, they possess inherent biophysical limitations. They reflect aggregate transsynaptic membrane potentials that are highly reference-and dipole geometry-dependent, lack cell-type specificity, are prone to electrical artifacts, and often arise from neural origins that remain difficult to identify with precision^1,2^.Currently, the most established method for population-level optical recording is fiber photometry using Genetically Encoded Calcium Indicators (GECIs)^3–5^. This technique uses thin (0.2–0.4 mm diameter) optical fibers to record from brain structures with minimal tissue damage^4,6^, but remains an indirect measure of neuronal voltage activity with slow temporal resolution The development of Genetically Encoded Voltage Indicators (GEVIs) has offered a way to bypass these constraints in vivo^7–14^. Unlike traditional electrodes, GEVIs are genetically targetable and possess the fast kinetics necessary to capture millisecond-scale membrane voltage changes and individual action potentials. Optimization of GEVIs now enables long-term recording at sufficient sensitivity to capture single spikes of dozens of neurons in behaving mice. Therefore, voltage imaging is a novel tool for a variety of neuroscience labs.Applying GEVIs to fiber photometry has been difficult because GEVI signals typically exhibit much lower SNR than their calcium counterparts^3,15,16^. Recent refinements in both sensor sensitivity and hardware optimization are beginning to change this. For instance, the “TEMPO” system^17,18^ and the recent uSMAART platform^8^ have demonstrated that GEVI-based fiber photometry is viable in awake, moving mice. These systems utilize sophisticated signal demodulation to extract robust voltage data and have shown that specific indicators, such as ASAP3, are particularly effective at capturing high-frequency subthreshold dynamics. Despite these advancements, several technical hurdles remain, particularly in terms of scalability and accessibility. Advanced photometry systems rely on specialized implementations using photomultiplier tubes (PMTs) or avalanche photodiode detectors (APDs) that are difficult to scale for simultaneous multi-site recordings^8,17^. In contrast, a sCMOS based imaging approach is already commonly used for cellular-level resolution GEVI imaging^7,10,13,19–21^ and the sCMOS pixel array would allow easier implementation of multi-site fiber photometry with GEVIs. Single-channel detectors, scaling them for simultaneous multi-site recordings requires duplicating expensive hardware for each additional site while sCMOS pixel arrays allow easier implementation by assigning regions of interest simultaneously for multi-site. Whereas commercially available fiber photometry systems have been optimized for GECIs (e.g. GCaMP) in terms of sensitivity and sampling rate, systems for voltage imaging are still under development. Widefield fluorescence microscopes are largely available in neuroscience institutes and a GEVI fiber photometry approach using standard widefield microscopes would significantly increase accessibility.

As noted by Legaria et al. (2022)^15^, GECI fiber photometry can be heavily contaminated by neuropil signals, meaning the recorded activity often reflects non-somatic changes rather than the direct output of the targeted cell bodies. Recent voltage fiber photometry studies^8,17^ use GEVIs that are expressed across the entire neuron, whereas cellular-resolution imaging often utilizes soma-targeted variants to reduce background fluorescence and increase signal-to-noise ratio^7,10,13,19,20,22^. It is currently unknown whether soma-targeted GEVIs can provide enough signal for robust population-level photometry. Transitioning to soma-targeted indicators would not only align cellular and population imaging protocols but also increase signal interpretability.

In this study, we address these challenges by demonstrating high-speed, multi-site fiber photometry GEVI dual-color imaging using a widefield sCMOS-based setup. This hardware configuration, originally designed for cellular-resolution voltage imaging, allows for the simultaneous recording of multiple fiber sites. By employing either the soma-targeted GEVI^21^ CamkII-pAce-KV2.1PR or CAG-DIO-AcemNeon2-KV2.1PR in awake, behaving mice, we capture high-fidelity voltage dynamics in a cell-type specific and soma-targeted manner. Our approach provides a scalable framework for monitoring cell-type-specific population GEVI across distributed neural circuits with high temporal precision.

## Results

### An integrated platform for simultaneous high-speed voltage imaging and electrophysiology

To bridge the gap between cell-type-specific voltage dynamics and traditional electrophysiology, we developed an integrated platform capable of high-SNR population voltage imaging using soma-targeted genetically encoded voltage indicators (GEVIs) and multi-site fiber photometry. Our optical configuration utilized a widefield microscope setup, optimized for cellular voltage imaging^20^, coupled with a high-speed sCMOS camera (Fig. 1A). Dual-wavelength excitation was achieved using a 488 nm laser for GEVI fluorescence and a 565 nm LED for anatomical localization and/or reference channel signal acquisition. To enable high-throughput data acquisition from distributed circuits, we employed a multi-fiber patch cable that projected the emission from multiple 200μm optical fibers onto discrete regions of interest (ROIs) on the sCMOS sensor. A digital mirror device was used to pattern the illumination to these restricted ROIs to optimize single-cell resolution voltage imaging^13,20,22^, but it is not necessary for fiber photometry population imaging. We first validated this platform in the hippocampal CA1 of awake, head-fixed mice on a treadmill, where neural oscillations are robustly modulated by locomotion^13,23–25^. To capture population-level voltage fluctuations, we expressed the soma-targeted GEVI, *pAce*, under the *CaMKII* promoter^27^ to target excitatory pyramidal neurons, while co-expressing *hSyn-mCherry* for structural reference. An implant configuration comprising a 200μm optical fiber attached with a twisted bipolar platinum-iridium wire for local field potential (LFP) recording was implanted into the CA1 (Fig. 1B). This dual-modal approach allowed for the simultaneous monitoring of cellular-level voltage transients and the extracellular LFP from the same neural substrate with the additional implantation of another LFP electrode on the contralateral hemisphere for recording LFP of the contralateral CA1 as well.

**Figure 1:**
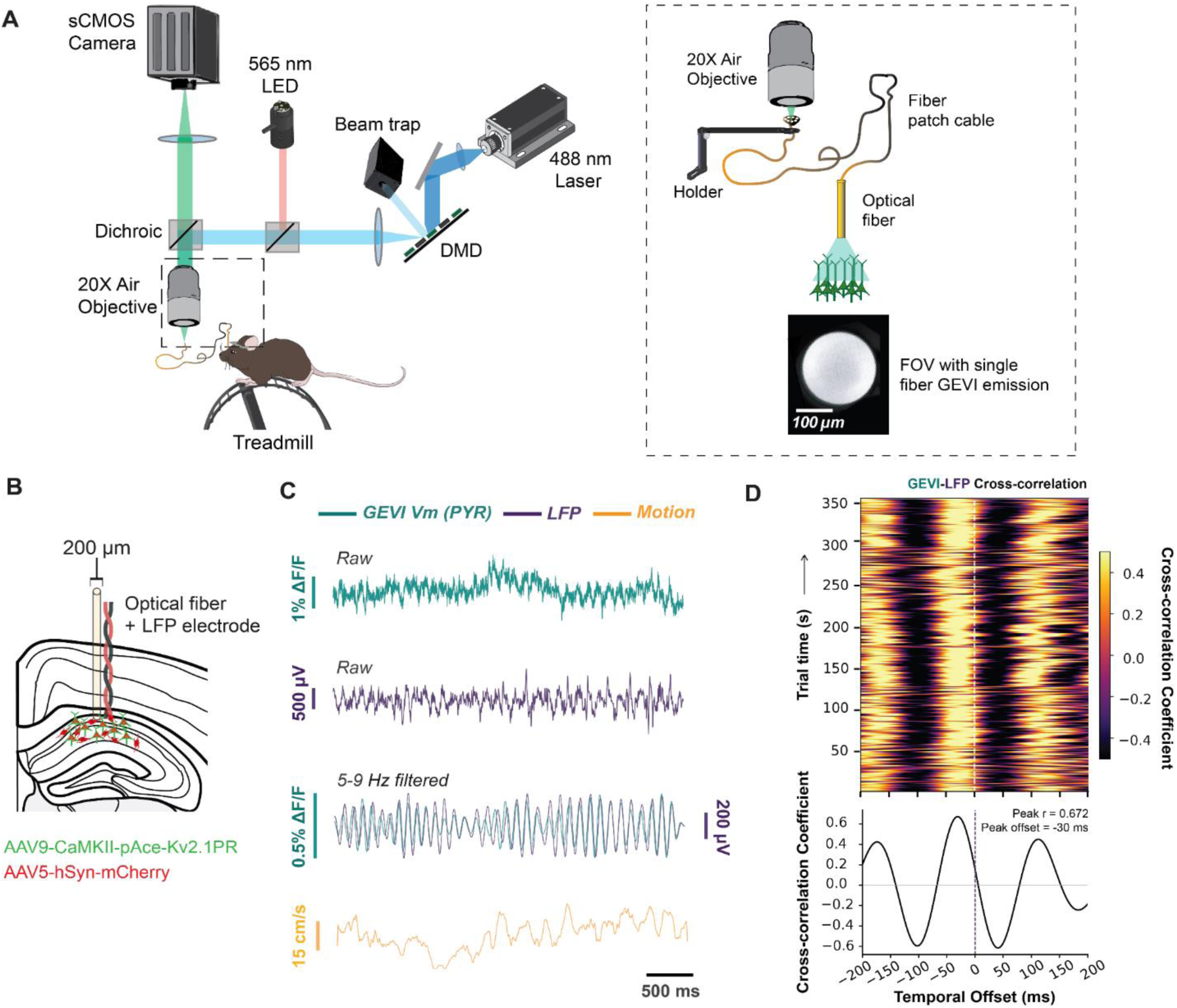
Integrated Platform for Deep-Brain Voltage Imaging and LFP Recording. (A, B) details the experimental configuration for fiber photometry recording in behaving mice. The optical path (**A**) uses a 488 nm laser and 565 nm LED for GEVI excitation and anatomical targeting, respectively. A patch cable enables simultaneous emission collection from two brain regions, as shown in the fiber FOV. Schematic **B** illustrates the 200 µm optical fiber and LFP electrode targeting hippocampal neurons expressing AAV9-CaMKII-pAce-Kv2.1PR (green) and AAV5-hSyn-mCherry (red). **C** Representative traces showing (top to bottom) GEVI fluorescence of CA1 pyramidal neurons, CA1 LFP, locomotion speed, and filtered rhythmic GEVI fluorescence and LFP. **D** Top illustration shows time-resolved cross-correlation between GEVI fluorescence and LFP signals, shown as a function of temporal offset (lag) and recording time. Warmer colors indicate stronger positive correlation. The bottom illustration shows the average cross-correlation function across the temporal offset if a 60s imaging trial.

Representative raw traces revealed that 498Hz sampled GEVI-derived fluorescence signals closely tracked LFP oscillations, particularly during periods of high-velocity locomotion (Fig. 1C). To quantify the temporal relationship between GEVI and LFP activity, we computed the cross-correlation between GEVI fluorescence and LFP signals. Time-resolved cross-correlation revealed a stable theta rhythmic oscillatory structure across the recording, indicating consistent coupling between the two modalities (Fig. 1D). Theta-band (5–9 Hz) cross-correlation between the GEVI signal and the LFP was robust and highly significant across all recording sessions (n = 21, 60 second imaging trials, across 5 separate recording days for one animal with median peak |r| = 0.673, mean peak lag = −19.5 ms [LFP leading]; 21/21 trials individually significant at p < 0.05; pooled p < 9.5 × 10⁻⁵; circular-shift surrogate test, 500 surrogates/trial), suggesting a systematic temporal offset between GEVI and LFP signals.

To assess the impact of behavioral state on neural dynamics, we compared spectral properties of LFP and GEVI CA1 signals during rest and running. Time–frequency analysis (Fig.2a) and power spectral density analysis (Fig.2b) revealed significantly increased narrow-band 6-8Hz theta frequency power during running in both LFP (Fig. 2B; p=0.0006) and GEVI signals (Fig. 2C; p=0.024). Notably, overall GEVI power prior to aperiodic correction was higher during rest compared to running, consistent with a previous report^13^. To assess whether GEVI signals reflect electrophysiological activity, we computed spectral coherence between GEVI and LFP signals. Theta-band coherence was significantly increased during running (Fig. 2D; Wilcoxon signed-rank test, one-tailed, p = 0.023, n = 6), indicating enhanced coupling between modalities. The GEVI signal quality, as quantified in terms of overall fluorescence (Suppl. Fig1a), over 6 x 1min-trial recordings showed an overall decline of around 1.5%. Within a trial, photobleaching rate exhibited a double exponential decay with initial faster decay and a slow sustained decay of 0.015% per sec (Suppl. Fig1B). Trials were separated by 10min inter-trial intervals. Fluorescence was recovered largely between trials. Signal quality assessed by GEVI-LFP CA1 theta coherence did not show decrease within and across trials (Suppl. Fig1C). Together, these results demonstrate that GEVI signals capture behaviorally modulated neural dynamics and remain tightly coupled to LFP activity.

**Figure 2:**
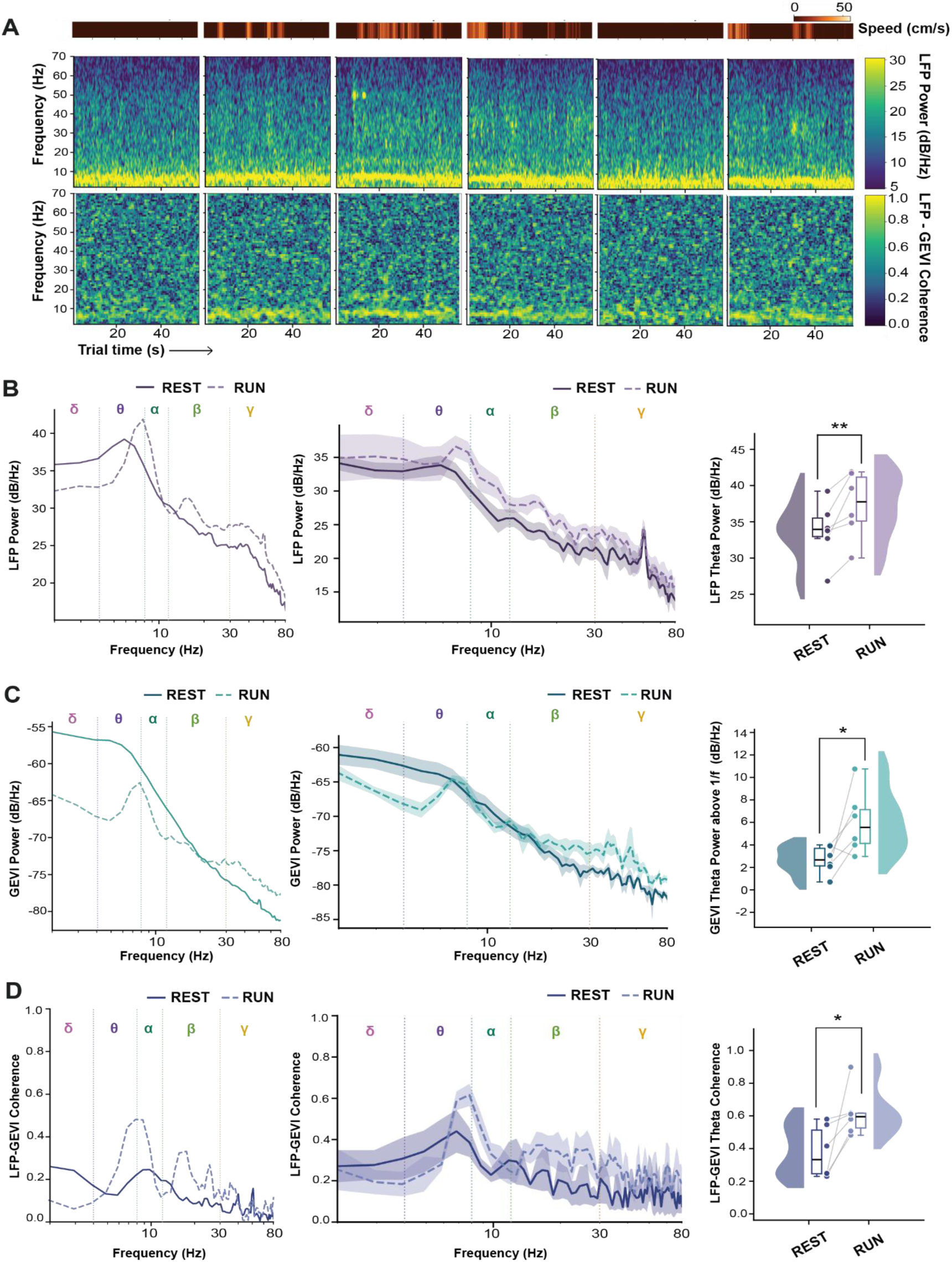
Locomotion-Dependent change in LFP and GEVI dynamics: Time-Frequency structure, spectral power and cross-signal coherence. (A) Speed heatmaps (top tow), time–frequency spectrograms of LFP (middle row) and LFP-GEVI coherence spectrograms (bottom row) across six representative 60s recording segments. Running epochs are associated with increased low-frequency power, particularly in the theta band. (B) LFP power spectral density (PSD) during rest and running. Example session (left column), population average across animals (middle column, mean ± s.e.m.) and quantification of band-limited theta power, showing a significant increase during running (right column). Normality of paired differences (rest-run) was assessed using one-tailed Wilcoxon signed-rank test (alternative = ‘greater’) was used, revealing a significant increase in theta power during running (p=0.006, n=6; one mouse was excluded due to insufficient rest data; effect size d=1.55). (C) GEVI power spectral density (aperiodic component removed, isolating oscillatory theta) during rest and running. Example session (left column), population average (middle column, mean ± s.e.m.), quantification of theta-band power (right column) showing increased GEVI power during locomotion (p=0.024, n=6, d=1.07). (D) Coherence between GEVI and LFP signals. Example session (left column), population average coherence spectra (middle; column, mean ± s.e.m.), summary quantification demonstrating increased GEVI–LFP coherence during running (right column; p=0.023, n=6, d=1.07).

### Pyramidal neuron specific Theta-Gamma phase-amplitude coupling in CA1 during locomotion states

A hallmark of hippocampal circuit function is the coordination of local gamma oscillations (30–80 Hz)^26,27^ by the phase of the theta rhythm (3–10 Hz), quantified as phase-amplitude coupling (PAC)^23,28,29^. Having established that our sCMOS-based photometry platform can resolve both theta -band CA1 dynamics, we next investigated whether this platform could detect cell-type-specific theta-gamma PAC in CA1 excitatory pyramidal neurons (Pyr) (Fig. 3).

**Figure 3:**
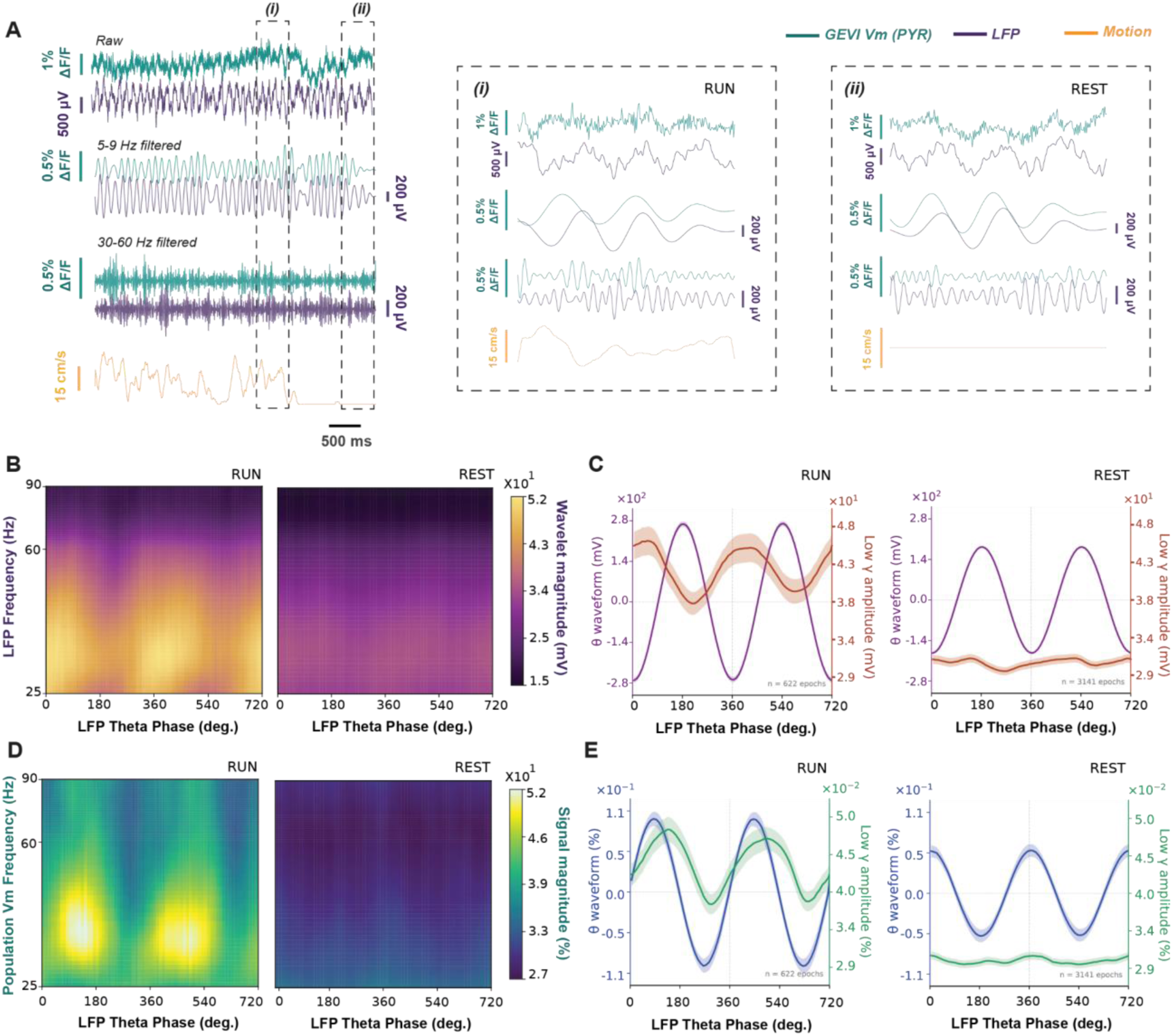
GEVI fiber photometry tracks theta-modulation of CA1 gamma modulation. (A-E) characterizes the cross-frequency coupling between LFP theta phase and gamma-band amplitude in Pyramidal (Pyr) neurons during rest and running states. (A) Representative traces showing (left column, top to bottom) raw GEVI fluorescence of Pyr neurons (blue), LFP (purple), bandpass-filtered signals, 5–9 Hz filtered activity and 30–60 Hz filtered activity, and motion (cm/s, orange). The second column (i) displays an example during a run period, while the third column (ii) shows an example trace during a rest epoch. (B) Phase-amplitude comodulograms for LFP signals during rest and running, showing wavelet amplitude as a function of LFP theta phase and frequency. Color indicates LFP wavelet magnitude. (C) Phase-resolved coupling for LFP signals. The theta-band waveform and low gamma-band amplitude are shown for the run (left) and rest (right) epochs. (D) Phase-amplitude comodulograms for GEVI fluorescence signals, plotted as in (B). Colour indicates fluorescence signal magnitude. (E) Phase resolved coupling for GEVI signals, plotted as in (C).

First, we evaluated the Pyr neuronal GEVI traces during rest and running time periods (Fig. 3A). We then calculated phase-amplitude comodulograms to visualize the distribution of gamma-band power relative to the LFP theta phase during rest and running. GEVI signals exhibited clear modulation of gamma amplitude by the underlying LFP theta phase (theta-gamma MI, running = 9.603×10⁻⁴, rest = 2.472×10⁻⁵, both p<0.001 based on surrogate testing, Fig.3 A,C) similar to LFP gamma-band signal (theta-gamma MI, running=5.414×10⁻⁴, rest=2.996×10⁻⁵, both p<0.001 based on surrogate testing, Fig. 3B, D).

The comodulograms during running showed a concentrated and higher-magnitude modulation of gamma power in both LFP signals and GEVI fluorescence (Fig. 3B, D). Specifically, the modulation depth of Pyr GEVI signal showed a tight phase-locking of gamma amplitude to the peak of the theta cycle, whereas LFP gamma amplitude was locked to the LFP theta through. Distinct phase-relation of GEVI gamma power to LFP theta phase in contrast to LFP gamma power has been also observed in a prior study^8^. This might be because of the more local and cellular specificity of the GEVI CA1 signal in contrast to LFP gamma.

These results demonstrate that GEVI fiber photometry not only captures the spectral components of the LFP but also resolves the intricate, cell-type-specific temporal cross-frequency relationships between different frequency bands. By revealing that Pyr populations exhibit distinct voltage-based PAC signatures, our platform provides a critical advantage over traditional LFP recordings, which represent a masked summation of all local currents.

### Population and single-cell dynamics during Deep Brain Stimulation

To evaluate the temporal resolution and sensitivity of our sCMOS-based platform under high-frequency perturbations, we performed electrical deep brain stimulation (DBS) in hippocampal CA1 while simultaneously recording GEVI-mediated voltage signals and LFPs from the stimulated locus. We compared two stimulation protocols: a 40 Hz “gamma-range” oscillation and a 135 Hz “high-frequency” oscillation, the latter being clinically relevant^30–32^. In fiber photometry mode, the GEVI signal (CaMKII-pAce-KV2.1) resolved individual stimulation-evoked voltage transients at 40 Hz (Fig. 4A). In contrast, during 135 Hz stimulation, the population voltage signal exhibited a sustained depolarizing shift (DC-like baseline increase) with superimposed high-frequency oscillations that were most prominent during the early stimulation period (Fig. 4B). Quantification of the GEVI signal (ΔF/F) across defined temporal epochs—transient (0–0.15 s), sustained (0.15–1 s), and post-stimulation (1–5.8 s)—revealed frequency-dependent dynamics (Fig. 4C, D). For 40 Hz stimulation, both transient and sustained periods showed significant increases relative to pre-stimulation baseline (transient: p = 2.26 × 10⁻⁶; sustained: p = 3.84 × 10⁻¹⁰), with the sustained response significantly exceeding the transient (p = 3.84 × 10⁻¹⁰), while post-stimulation activity returned to baseline (p = 0.557). For 135 Hz stimulation, all epochs differed significantly from each other (Friedman test p = 2.33 × 10⁻⁶; all pairwise comparisons p = 0.012), indicating a robust and sustained depolarization that persisted into the post-stimulation period. Direct comparison between stimulation frequencies confirmed distinct response profiles (Fig. 4E). Within-frequency analyses showed that sustained responses exceeded transient responses for both 40 Hz (paired t-test, p = 2.56 × 10⁻¹⁰) and 135 Hz stimulation (Wilcoxon, p = 1.95 × 10⁻³). Between frequencies, 135 Hz stimulation elicited significantly larger ΔF/F responses than 40 Hz during both transient (p = 8.79 × 10⁻⁴) and sustained periods (p = 7.38 × 10⁻⁴). Time–frequency analysis demonstrated that the optical signal reliably tracked the DBS-induced effects (Fig. 4F, G), with clear power increases centered at 40 Hz and 135 Hz stimulation frequencies. Quantification of stimulation-band power (35–45 Hz for 40 Hz DBS; 130–140 Hz for 135 Hz DBS) revealed significant increases during sustained compared to transient periods for both frequencies (40 Hz: p = 1.95 × 10⁻³; 135 Hz: p = 2.27 × 10⁻⁷), with stronger overall power observed for 135 Hz stimulation (transient: p = 2.85 × 10⁻¹⁵; sustained: p = 3.65 × 10⁻⁴; Fig. 4H). We observed a similar 40Hz entrainment and depolarization in the striatum (Supplementary figure 2).

**Figure 4:**
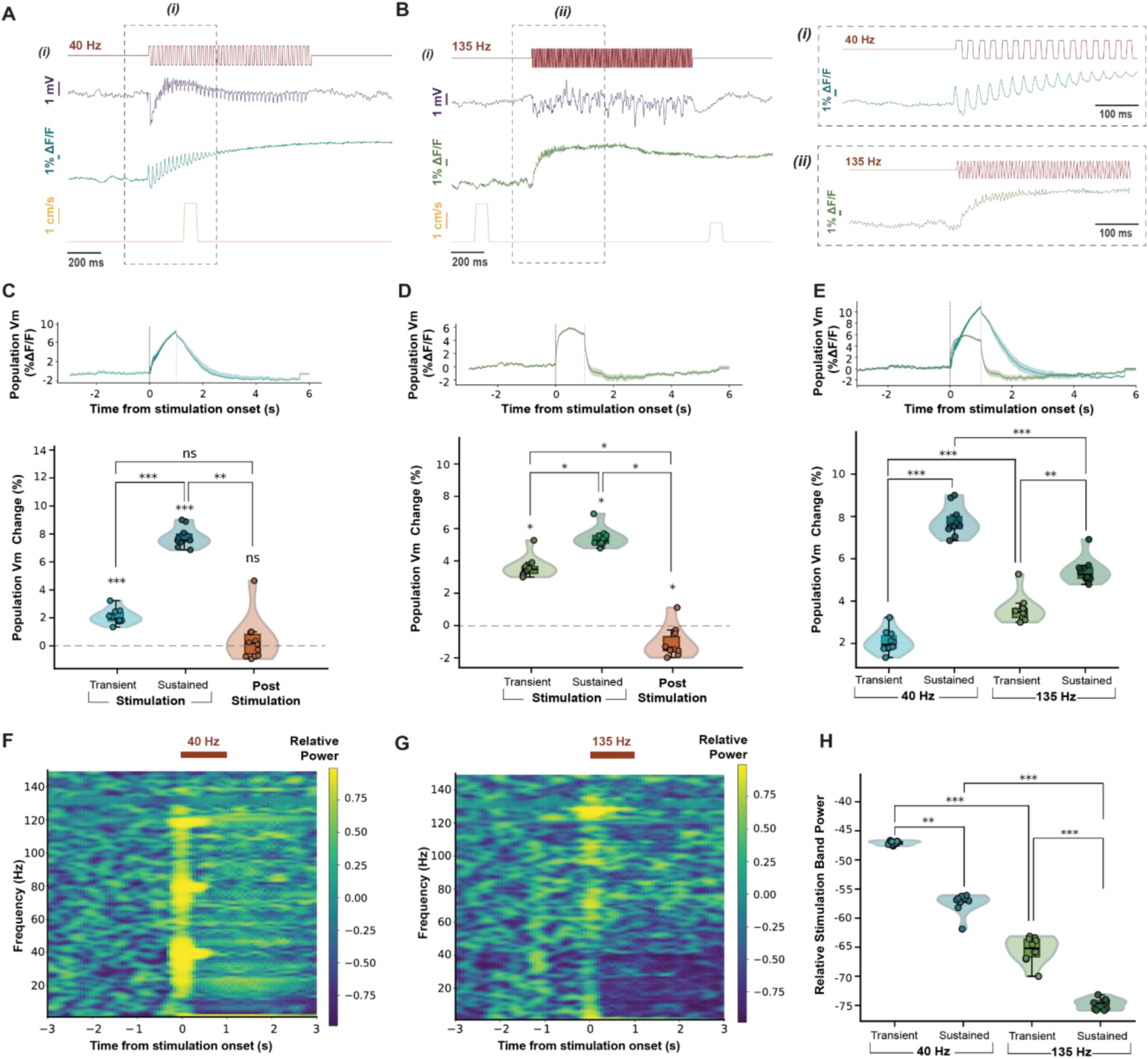
GEVI fiber photometry captures DBS neural modulation. (A, B) Representative single trial recordings showing LFP (top, purple), GEVI fluorescence (middle, teal), and movement artifacts signals (bottom, orange) during different 40Hz DBS (A) and 135 Hz stimulation (B, left). Insets (i-ii) show zoomed views of the same traces of 40Hz (i) and 135Hz (ii; B, right). top bars indicate stimulation periods. (C, D) Quantification of GEVI fiber Vm signal relative to baseline for transient (0-0.15s), sustained (0.15-1s) and post-stim (1-5.8s) periods during energy-balanced DBS. Average GEVI fluorescence traces (mean ± s.e.m; top), Violin plots showing distributions across trials (bottom). Statistical significance was assessed using a Friedman non-parametric repeated-measures test followed by pairwise post-hoc comparisons with Holm-Bonferroni correction. For 40 Hz DBS (C): Friedman χ²(3) = 25.56, p = 1.18 × 10⁻⁵, n = 10. Post-hoc (Holm-corrected): pre vs transient, paired t-test, df = 9, p = 2.26 × 10⁻⁶; pre vs sustained, paired t-test, df = 9, p = 3.84 × 10⁻¹⁰; pre vs post, Wilcoxon, p = 0.557; transient vs sustained, paired t-test, p = 3.84 × 10⁻¹⁰; transient vs post, Wilcoxon, p = 0.074; sustained vs post, Wilcoxon, p = 5.86 × 10⁻³. For 135 Hz DBS (D): Friedman χ²(3) = 28.92, p = 2.33 × 10⁻⁶, n = 10. Post-hoc (Holm-corrected; Wilcoxon signed-rank for all comparisons): pre vs transient, p = 0.012; pre vs sustained, p = 0.012; pre vs post, p = 0.012; transient vs sustained, p = 0.012; transient vs post, p = 0.012; sustained vs post, p = 0.012. (E) Comparison of Vm responses between 40Hz and 135Hz DBS. Population-averaged traces (top), and violin plots comparing transient and sustained responses (bottom). Within-frequency comparisons (transient vs sustained) are paired; between-frequency comparisons are unpaired. Holm-Bonferroni correction was applied. For 40 Hz: paired t-test, df = 9, p = 2.56 × 10⁻¹⁰.For 135 Hz: Wilcoxon signed-rank, p = 1.95 × 10⁻³. Between frequencies: transient, Mann–Whitney U, p = 8.79 × 10⁻⁴; sustained, Mann–Whitney U, p = 7.38 × 10⁻⁴. (F-G) Time-frequency spectrogram during DBS for 40Hz (F) and 135Hz (G) conditions, computed using a 0.6s sliding window with 88% overlap, and shown as relative power changes from baseline. (H) Fiber stimulation-band power (dB relative to baseline; 40Hz: 35-45Hz band, 135Hz: 130-140Hz band) across transient and sustained periods. For 40 Hz: Wilcoxon signed-rank, p = 1.95 × 10⁻³. For 135 Hz: paired t-test, df = 9, p = 2.27 × 10⁻⁷. Between frequencies: transient, independent t-test, df = 18, p = 2.85 × 10⁻¹⁵; sustained, Mann–Whitney U, p = 3.65 × 10⁻⁴.

A unique advantage of our sCMOS widefield approach is the ability to transition from population-level fiber photometry to cellular-resolution imaging by modifying the optical interface. We leveraged this to investigate the recruitment of individual CA1 neurons during DBS (Fig. 5) using CaMKII-pAce-KV1.2. Using DMD-based selection of soma-restricted ROIs, we identified and recorded from 11 distinct neurons (N1–N11) (Fig. 5A, left).

**Figure 5:**
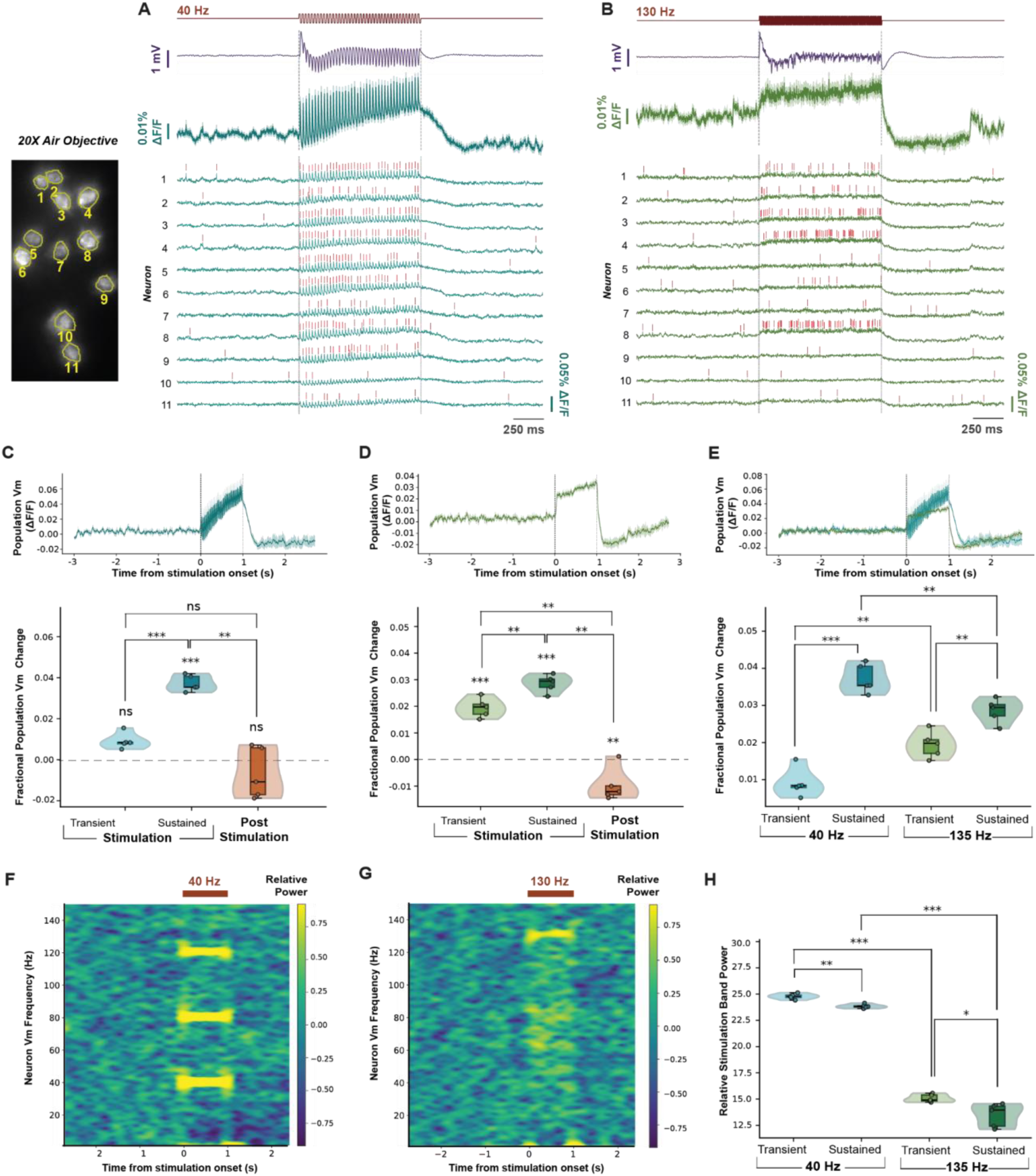
GEVI single-cell recordings capture DBS neural modulation. (A, B) Simultaneous pAce-KV2.1 voltage imaging and LFP recordings during 40 Hz (A) and 130 Hz (B) DBS. Left: field of view (FOV) with segmented neurons (N1–N11) and corresponding masks. Middle/right: representative single-trial recordings, showing population-averaged Vm signal across all neurons (top, green), individual neuron traces (bottom, green), LFP (top, purple), and stimulation periods (shaded). Red ticks indicate detected spikes. (C, D) Quantification of population Vm signal (ΔF/F; averaged across 11 neurons per trial) across pre-stim (−3–0 s), transient (0–0.15 s), sustained (0.15–1 s), and post-stim (1–3 s) periods during energy-balanced DBS. Top: population-averaged Vm traces (mean ± s.e.m.). Bottom: violin plots showing trial distributions (n = 5 trials), with kernel density (outer shape), interquartile range (box; 1× and 1.5×), median (black line), and individual trials (scatter points). Statistical analysis used a Friedman non-parametric repeated-measures test followed by Holm–Bonferroni-corrected pairwise comparisons (paired t-test, based on Shapiro–Wilk normality testing; 6 comparisons). For 40 Hz DBS (C): Friedman χ² (3) = 14.04, p = 2.85 × 10⁻³, n = 5. Post-hoc (paired t-test, df = 4): pre vs transient, p = 0.085; pre vs sustained, p = 4.96 × 10⁻⁵; pre vs post, p = 0.094; transient vs sustained, p = 7.57 × 10⁻⁵; transient vs post, p = 0.085; sustained vs post, p = 1.49 × 10⁻³. For 130 Hz DBS (D): Friedman χ²(3) = 15.00, p = 1.82 × 10⁻³, n = 5. Post-hoc (paired t-test, df = 4): pre vs transient, p = 5.86 × 10⁻⁴; pre vs sustained, p = 3.02 × 10⁻⁴; pre vs post, p = 9.62 × 10⁻³; transient vs sustained, p = 9.62 × 10⁻³; transient vs post, p = 2.07 × 10⁻³; sustained vs post, p = 1.22 × 10⁻³. (E) Comparison of transient and sustained Vm responses between 40 Hz and 130 Hz DBS. Top: population-averaged Vm traces. Bottom: violin plots comparing conditions (40 Hz: n = 5 trials; 130 Hz: n = 5 trials; 11 neurons averaged per trial). Within-frequency comparisons (transient vs sustained) are paired; between-frequency comparisons are unpaired. Holm–Bonferroni correction was applied (4 comparisons). For 40 Hz: paired t-test, df = 4, p = 6.06 × 10⁻⁵. For 130 Hz: paired t-test, df = 4, p = 9.62 × 10⁻³. Between frequencies: transient, independent t-test, df = 8, p = 6.58 × 10⁻³; sustained, independent t-test, df = 8, p = 9.62 × 10⁻³. (F, G) Time–frequency representations (TFRs) of population Vm signals during DBS for 40 Hz (F) and 130 Hz (G), computed using a 0.6 s sliding window with 95% overlap, shown as relative power changes from pre-stimulation baseline. (H) Population Vm stimulation-band power (dB relative to pre-stim baseline; 40 Hz: 35–45 Hz band; 130 Hz: 125–135 Hz band) during transient and sustained periods. Holm–Bonferroni correction applied (4 comparisons). For 40 Hz: paired t-test, df = 4, p = 5.32 × 10⁻³. For 130 Hz: paired t-test, df = 4, p = 0.024. Between frequencies: transient, independent t-test, df = 8, p = 1.52 × 10⁻¹⁰; sustained, independent t-test, df = 8, p = 1.02 × 10⁻⁷.

At 40 Hz, single-trial recordings revealed highly structured, pulse-locked spiking and subthreshold oscillations across the majority of sampled neurons (Fig. 5A). In contrast, 130 Hz stimulation evoked a sustained, plateau-like depolarization in somatic voltage across neurons, with reduced temporal locking to individual stimulation pulses (Fig. 5B). Quantification of the population-averaged single-cell Vm signal (ΔF/F; 11 neurons averaged per trial) across defined temporal epochs—transient (0–0.15 s), sustained (0.15–1 s), and post-stimulation (1–3 s)—revealed distinct frequency-dependent dynamics (Fig. 5C, D). For 40 Hz stimulation, the sustained response was significantly elevated relative to baseline (p = 4.96 × 10⁻⁵), while the transient response did not reach significance (p = 0.085). The sustained response was also significantly greater than the transient (p = 7.57 × 10⁻⁵), and activity returned toward baseline post-stimulation. In contrast, 130 Hz stimulation induced significant increases across all epochs compared to baseline (transient: p = 5.86 × 10⁻⁴; sustained: p = 3.02 × 10⁻⁴; post-stim: p = 9.62 × 10⁻³), with all pairwise comparisons reaching significance, consistent with a robust and persistent depolarization.

Direct comparison between stimulation frequencies confirmed these differences (Fig. 5E). Within-frequency analyses showed that sustained responses exceeded transient responses for both 40 Hz (paired t-test, p = 6.06 × 10⁻⁵) and 130 Hz stimulation (paired t-test, p = 9.62 × 10⁻³). Between frequencies, 130 Hz stimulation elicited significantly larger responses than 40 Hz during both transient (p = 6.58 × 10⁻³) and sustained periods (p = 9.62 × 10⁻³). Time–frequency analysis of the population-averaged single-cell signals demonstrated clear entrainment of the optical signal to the stimulation frequency (Fig. 5F, G). Quantification of stimulation-band power (40 Hz: 35–45 Hz; 130 Hz: 125–135 Hz) revealed significant increases during the sustained compared to transient period for both frequencies (40 Hz: p = 5.32 × 10⁻³; 130 Hz: p = 0.024), with substantially greater power observed during 130 Hz stimulation (transient: p = 1.52 × 10⁻¹⁰; sustained: p = 1.02 × 10⁻⁷; Fig. 5H).

Importantly, these cellular-resolution measurements closely recapitulate the dynamics observed at the population level using fiber photometry (Fig. 4) as well as in line with a prior cellular GEVI study^32^, demonstrating that the apparent transition from temporally precise, pulse-locked activity at 40 Hz to sustained depolarization at higher frequencies emerges consistently across spatial scales. Collectively, these results highlight the versatility of the sCMOS-based system to capture high-speed neural dynamics ranging from single-neuron activity to population-level signals, providing a powerful framework for dissecting the circuit-level effects of neuromodulation.

### Dual-site GEVI-LFP recordings reveal locomotion-dependent enhancement of interhemispheric CA1 theta coupling

To examine large-scale neural coordination during behavior, we first performed simultaneous bilateral 498Hz recordings of GEVI fluorescence and LFP signals in hippocampal CA1. This approach enabled direct comparison of neural dynamics across spatially distinct recording sites within the same animal (Fig. 6A). Representative recordings from left and right CA1 (Fig. 6B, C) demonstrate temporally aligned GEVI and LFP activity across hemispheres. Locomotion epochs were associated with pronounced rhythmic activity in both regions, particularly in the theta frequency range (5–9 Hz), as observed in band-pass filtered traces.

**Figure 6:**
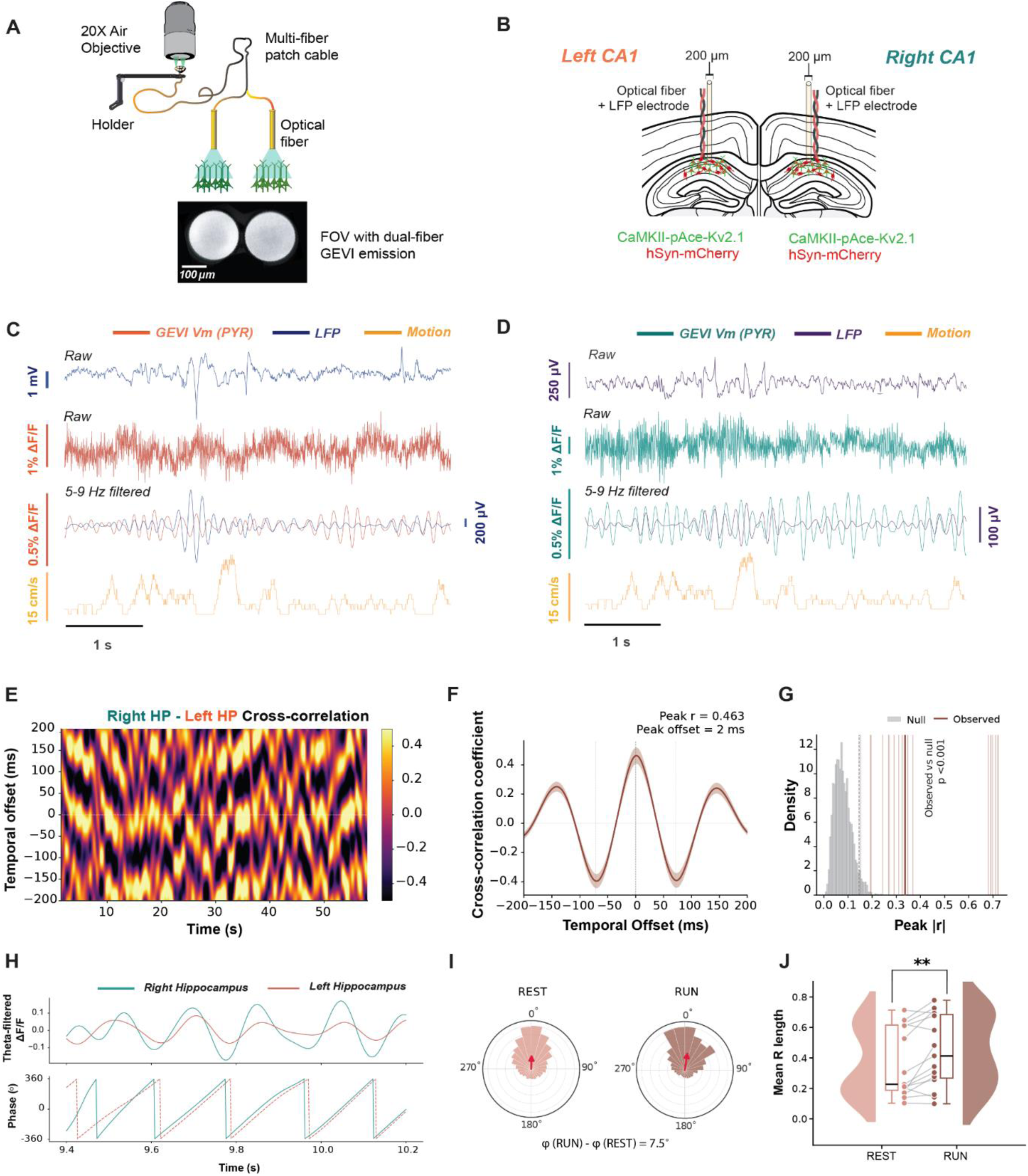
Multi-site GEVI and LFP recordings enable simultaneous characterization of spectral dynamics and interhemispheric coordination during locomotion. (**A**) The dual-fiber patch cable enables simultaneous emission collection from two brain regions, as shown in the fiber FOV captured by the sCMOS camera. (**B**) Schematic of the experimental configuration showing bilateral CA1 recordings, optical fiber placement, LFP electrodes, and viral constructs (CaMKII–pAce-Kv2.PR1 and AAV5-hSyn-mCherry). (**C, D**) Representative simultaneous recordings from left (**C**) and right (**D**) CA1 (5 s excerpt), showing LFP (top), GEVI fluorescence (ΔF/F; middle), theta band-pass filtered signals (5–9 Hz), and locomotion speed (bottom). Locomotion epochs are associated with increased theta-band rhythmicity in both hemispheres. (**E**) Time-resolved cross-correlation heatmap for an example trial (including both rest and locomotion epochs), illustrating dynamic inter-hemispheric coupling over time. (**F**) Average cross-correlation function across trials showing a clear peak (r = 0.463) at a lag of +2 ms (positive lag indicates right hippocampus leading), consistent with temporally structured coordination (n = 18 trials, 1 mouse). (**G**) Circular shift surrogate analysis (n = 500 surrogates per trial) validating cross-correlation strength. Grey bars indicate the null distribution (circularly shifted data), the thick brown line indicates the observed median |r| = 0.336, and the dashed black line marks the 95th percentile of the null distribution (|r| = 0.146). Observed correlations significantly exceed the null distribution (pooled median vs null, p < 0.001). (**H**) Theta-filtered GEVI signals (ΔF/F; top) and corresponding instantaneous Hilbert phase (bottom) from bilateral CA1 during a representative ∼1 s recording segment, illustrating phase-locked oscillatory activity across hemispheres. (**I**) Polar (rose) plots of phase differences between hemispheres during rest and locomotion, demonstrating a non-uniform phase distribution with stronger clustering during locomotion. (**J**) Quantification of phase-locking strength using mean resultant vector length (R) per trial. Locomotion significantly increases phase synchronization compared to rest (paired t-test, p = 0.0070, n = 15 trials; mean R_run = 0.443, mean R_rest = 0.365).

To quantify inter-hemispheric coordination, we computed time-resolved cross-correlations between GEVI signals. Cross-correlation heatmaps (Fig. 6D) revealed dynamic coupling across both rest and locomotion periods. The average cross-correlation function exhibited a clear peak (r = 0.463) at a lag of +2 ms, indicating that activity in the right hippocampus slightly leads the left (Fig. 6E; n = 18 trials). To assess statistical significance, we performed a circular shift surrogate analysis (500 surrogates per trial). The observed median correlation magnitude (|r| = 0.336) exceeded the 95th percentile of the null distribution (|r| = 0.146), confirming that inter-site coupling was significantly greater than expected by chance (p < 0.001; Fig. 6F). Consistent bilateral hippocampal theta-band (5–9 Hz) synchrony was also confirmed in a second animal (n = 8 60 second imaging trials across 2 recording days): the right HP - left HP multi-site fiber cross-correlation yielded a median per-trial peak |r| = 0.787 at a mean peak lag of 0.5ms; all 8/8 trials were individually significant (p < 0.05), pooled p < 9.5 × 10⁻⁵ (circular-shift surrogate test, 500 surrogates/trial).

We next examined phase relationships between hemispheres using theta-filtered signals. Phase-resolved traces (Fig. 6G) demonstrated coherent oscillatory structure across sites in line with prior reports^33,34^. Polar histograms of phase differences (Fig. 6H) revealed a non-uniform distribution, indicating a preferred phase relationship that became more pronounced during locomotion. Quantification of phase-locking strength using the mean resultant vector length (R) showed a significant increase during locomotion compared to rest (paired t-test, p = 0.0070, n = 15 trials; mean R_run = 0.443 vs mean R_rest = 0.365; Fig. 6I), indicating enhanced phase synchronization across hemispheres during active behavior.

Together, these results demonstrate that simultaneous bilateral GEVI recordings capture coordinated, lag-structured, and phase-locked activity across hippocampal networks. Locomotion not only enhances local theta oscillations but also strengthens long-range inter-hemispheric coupling and phase synchronization, highlighting the ability of this platform to resolve distributed circuit dynamics.

### Dual-color, dual-site fiber GEVI imaging simultaneously in multiple freely-running mice

Mechanical and biological factors can induce substantial temporal variability of signal quality. Typical artefacts are mechanical forces on the patch cable or fiber cannula that modulates transmission, particularly in free-running case, and biological non-neuronal sources like hemodynamic changes. Interpretability of fluorescence signals can be improved if a control fluorophore is simultaneously recorded as a reference signal. Control fluorophore is a static fluorescence protein that is not voltage-dependent and is typically in a non-overlapping wavelength, e.g. red-shifted when recording blue-green GEVIs. Dual-color imaging with sCMOS is not straightforward and can be cost-intensive if two sCMOS are used for each wavelength^8,21^. Here, we employ a single sCMOS solution for dual color imaging by splitting the emission by color and projecting them onto distinct FOVs on the sCMOS camera (Figure 7a). The emission is first demagnified by the tube lens (0.6x) and then split by a dichroic mirror. With the use of two adjustable mirrors, the green and red emission are projected onto separate nearby FOVs on the sCMOS. All components for emission color splitting are commercially available and easy to assemble. To demonstrate the feasibility of the approach, we imaged a 5-core fiber bundle in dual-color in three freely-running PV-cre mice injected with CAG-DIO-AcemNeon2-KV2.1PR and hSyn-mCherry in dorsal CA1 and medial prefrontal cortex (mPFC) at 430Hz. In 2 PV-cre mice, we recorded CA1 and mPFC simultaneously as well as from one additional CA1 from the third PV-cre mouse (see Figure 7b). Like previous experiments, we used 40Hz DBS 1-sec stimulation in CA1 to assess neuronal entrainment in the PV interneurons of the three mice. We found clear 40Hz DBS entrainment, reflected in specific 40Hz spectral power increase, in PV interneuron signals from CA1s in all three mice and weaker 40Hz entrainment in PV interneurons from mPFCs (Figure 7E,G). In contrast, no 40Hz entrainment was observed in control fluorophore mCherry demonstrating that the DBS effects are somatic membrane voltage dependent (Figure 7F,H). Overall, these results demonstrate the feasibility of sensitive multi-fiber, dual-color high-speed fiber voltage photometry in freely-running mice.

**Figure 7:**
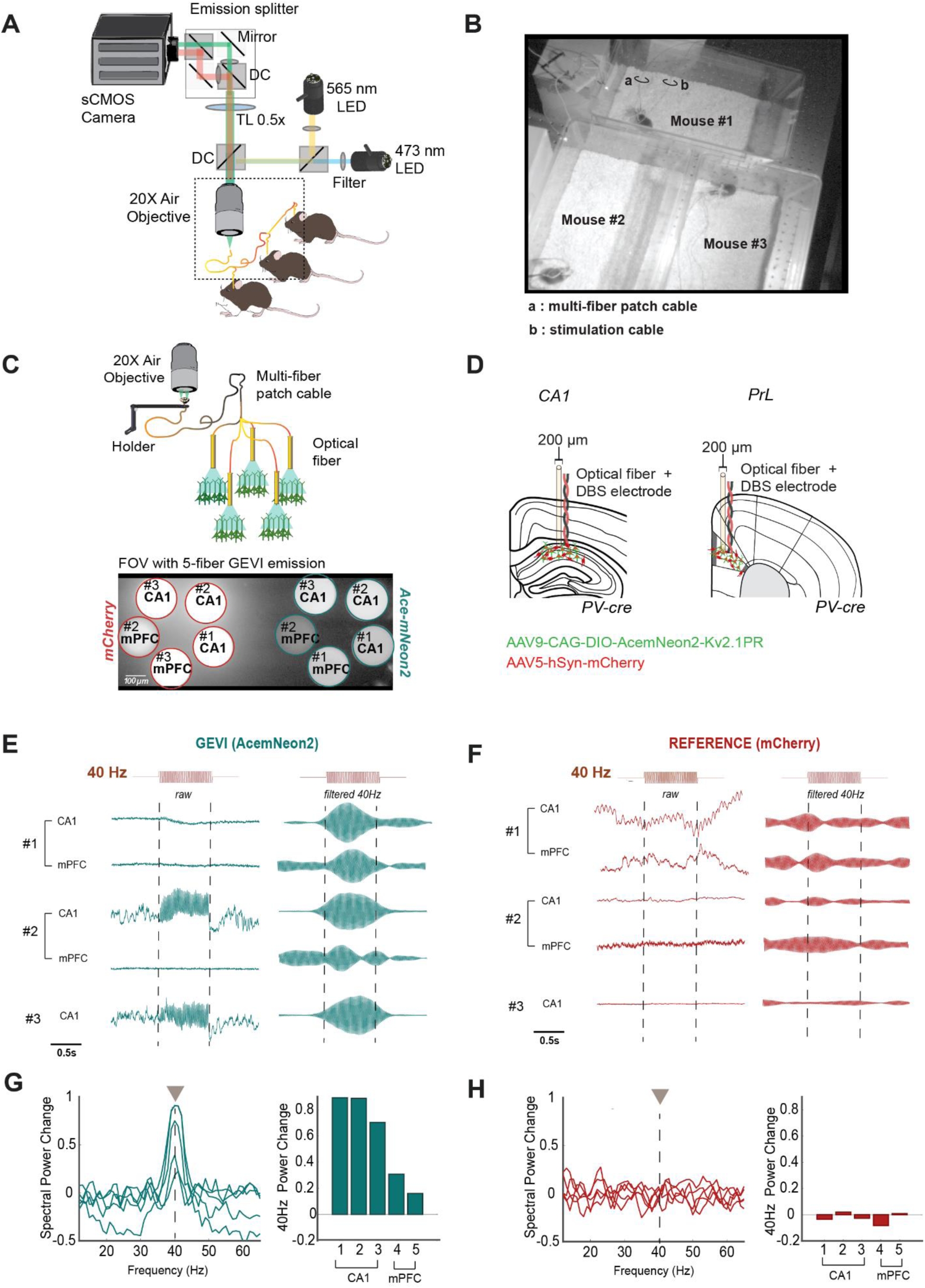
High-throughput dual-color fiber voltage imaging in freely running mice during DBS. **(A)** Schematic overview of the 2^nd^ microscope setup that includes emission beam splitting for dual color voltage imaging in the blue (470nm) and yellow (LED, 565nm) for AcemNeon2 and mCherry respectively. (**B**) Using a 5-fiber patch cable, we recorded in three PV-cre mice simultaneously. (**C**) At the level of sCMOS, the FOV of the 5-fiber surfaces is represented for green emission (AcemNeon2) and mirrored for red emission (mCherry).(**D**) 2 PV-cre animals had fiber with electrodes in CA1 and mPFC and 1 PV-cre mice only in CA1. (**E**) The recorded GEVI signals, left raw and right filtered at 40Hz, during 1sec 40Hz electrical stimulation in CA1. Red trace (top) represents the electric stimulations pulses. (**F**) Same as (E), but for simultaneously recorded mCherry signals. (**G**) The spectral power change relative to baseline for the 5 simultaneously recorded GEVI signals. An increase in spectral power at 40Hz during 40Hz electrical stimulation suggests entrainment. Quantification of 40Hz spectral peak is shown on the right. Strong entrainment is seen in CA1. (**H**) Same as (**G**), but for mCherry. No spectral power changes in mCherry signals were observed at 40Hz during 40Hz electrical stimulation.

## Discussion

In this study, we show that a widefield microscope equipped with a high-speed sCMOS camera can be adapted into a scalable, multi-site GEVI fiber photometry platform. Using soma-targeted GEVIs, we obtained high-SNR recordings of both somatic membrane oscillations and high-frequency transients across distributed circuits, including the bilateral hippocampus CA1 subfields, medial prefrontal cortex and the striatum. We demonstrate compatibility of our optical platform with simultaneous recording of classical electrophysiological measurements such as LFP and robustly capture behaviorally relevant dynamics and high-frequency electric stimulation effects in pyramidal and interneuron populations. GEVI photometry resolved baseline CA1 dynamics with temporal precision comparable to LFP capturing theta (3–10 Hz) and gamma (30–80 Hz) as well as theta–gamma cross-frequency coupling. We also show that this platform spans spatial scales, from population-level fiber photometry to single-cell resolution voltage imaging. By extending the recordings across multiple sites, hemispheres and animals, we demonstrate the high throughput capacity of the imaging system enabling future multi-area and multi-animal recordings.

### Scalable multi-site GEVI monitoring

Our sCMOS-based approach offers inherent scalability. Traditional photometry systems require a separate photodetector (PMT or photodiode) for each recording site, which becomes increasingly complex as the number of sites increases^8^. By treating each optical fiber as a region of interest on a single large-scale sensor, our platform can image multiple fibers simultaneously without the need for additional cameras. The high data throughput of the sCMOS enables studies of inter-regional and inter-animal voltage dynamics, such as the synchronized bilateral CA1 activity or multi-animal imaging demonstrated here. Usage of sCMOS technology for fiber photometry has been used for multi-site fiber calcium photometry and several commercial devices are available using this technology for calcium fiber photometry.

### Efficient dual color GEVI fiber imaging

Dual-color imaging for fiber photometry is often necessary to assess with a control fluorophore whether observed signal fluctuations are from the relevant cell population or due to artefacts from movement, hemodynamics or other processes^16,35,36^. To enable dual-color sCMOS imaging, integrating two sCMOS cameras for each wavelength has been demonstrated for population voltage imaging^8^. However, due to the costs of a high-speed sCMOS and the amount of data created, this is not a cost-effective and data-effective implementation. We therefore opted for a one sCMOS camera solution, in which the FOVs of each wavelength are projected not on two sCMOS, but onto the same sCMOS. This was enabled by using a demagnifying tube lens (0.6x) and by splitting the emission based on wavelength and projecting the FOVs next to each other on the sCMOS sensor. This allowed simultaneous dual-color imaging in freely running mice (Fig.7).

#### Direct somatic voltage imaging

Prior voltage fiber photometry^8,17,18^ have used GEVI expression across the whole neurons’ membrane, which is also common for other indicators like gCaMP^16,37^. However, the biophysical interpretation of neuronal signaling is complicated, because e.g. dendritic or somatic signaling contribute differently to the measured signal^15^. While fiber photometry studies have used GEVIs without specific targeting, cellular voltage imaging studies commonly use soma-targeted GEVIs to reduce background fluorescence, which would mean that labs need distinct GEVI indicators for both recording modalities. Here, we show that robust fiber photometry GEVI imaging, comparable to prior studies^8,17,18^, can be obtained using soma-targeted GEVIs. Soma-targeting thus enables capturing the primary output of the targeted population and aligning population-level photometry with cellular-resolution measurements with the same GEVI and microscope setup. While the GEVI used is sensitive to subthreshold membrane voltage (see Fig 4 and 5), it is likely that soma-targeted GEVI fiber photometry has significant contribution by suprathreshold spiking activity. Comparison of GEVI fiber photometry with cellular-level voltage imaging can address this question. Future GEVI development might provide novel GEVIs with selective sensitivity to subthreshold or suprathreshold membrane voltage changes. Further, GEVI expression in other neuron compartments, e.g. axons, will provide additional understanding of the critical membrane voltage dynamics.

### High-frequency temporal sensitivity

Voltage imaging, similar to electrophysiology, demands for higher temporal resolution than calcium imaging. Our system captures theta (4-8Hz) and gamma rhythms (30-90Hz) as well as >100 Hz responses during electrical stimulation, demonstrating the temporal resolution and low read noise of the sCMOS sensor. Unlike GECIs, which act as low-pass filters due to the slower rise and decay time constants, GEVI photometry resolves sub-millisecond depolarization and repolarization kinetics, providing a reference-free optical analog of the LFP with higher cellular and spatial specificity. The sampling rate, here approximately 0.5kHz-2kHz, is limited by the number of lines the sCMOS needs to record to capture all the fiber outputs and the overall signal-to-noise (SNR), which is expected to decrease with sampling rate due to increased read-out noise. This high-frequency sensitivity enables the detection of rapid entrainment to external rhythms, as well as precise tracking of phase-dependent dynamics such as theta-gamma coupling^23,38^ and other cross-frequency interactions^28^. Moreover, the ability to resolve subthreshold fluctuations allows direct observation of membrane potential oscillations that shape neuronal excitability but are largely inaccessible with calcium-based approaches. Together these features enable a more complete characterization of circuit dynamics, linking fast synaptic activity, oscillatory coordination and population level signaling within the same measurement framework.

#### Comparison to previous voltage fiber photometry

Previous fiber photometry systems for GEVI imaging^8,17,18^ have used avalanche photodetectors (APD) to capture GEVI signals using non-soma-targeted GEVIs. Further, they implemented phase-sensitive illumination laser modulation and detection to increase SNR of GEVI signals. Here, the fiber photometry system is based on a sCMOS-based widefield imaging setup that has been originally optimized for cellular voltage imaging using soma-targeted GEVIs. sCMOS have high quantum efficiency (QE) and can record multiple fibers simultaneously, while APDs have high sensitivity (low read out noise) in low photon count conditions, requiring lower illumination conditions and are very fast as a single-pixel detectors allowing for phase-sensitive demodulation. Here, we choose a high-speed sensitive sCMOS as our approach, because it allows (1) recording from multiple fibers and colors simultaneously, (2) allows switching from population to single-cell resolution mode, (3) provides high SNR imaging of GEVI signaling in of pyramidal and interneuron somatic membrane voltage signals in head-fixed and freely running mice.

### Limitations

Despite these advantages, certain limitations exist. High-speed acquisition generates massive datasets at GB/s rates, requiring substantial computational resources for real-time streaming and post hoc analysis. After fiber trace extraction, the raw imaging files can however be deleted or compressed. Although the sCMOS quantum efficiency exceeds 90%, SNR is still limited by the brightness of current GEVIs relative to calcium indicators, and further improvements in fluorophore photostability and brightness will be needed for long-term or deep-tissue imaging. Finally, while soma-targeting improves signal specificity, fiber photometry remains a bulk measurement, averaging activity across all expressing neurons within the collection volume. Future integration of scalable sCMOS imaging with high-density fiber bundles may enable multi-site, single-cell resolution.

## Methods

### First microscope setup

A high-speed sCMOS camera (Teledyne, Kinetix) was optimized for high-speed, low-light imaging applications, making it well-suited for widefield voltage imaging (>90% peak quantum efficiency, as low as 0.7 e⁻), enabling high sensitivity, image quality, and data throughput (GB/s), and allowing recordings of large fields of view at millisecond resolution (up to 5kHz).

A continuous-wave (CW) diode laser emitting at 488 ± 5 nm acts as *excitation illumination source* for widefield voltage imaging. The illumination irradiance range was 3-5 mW/mm^2^. A multimode fiber collimator (FC/PC connector, Thorlabs F950FC-A) is used to collimate light from a high-NA multimode fiber.

Excitation beam attenuation and shuttering is achieved using an optical beam shutter (Ø 1″ aperture, e.g. Thorlabs SH1/M). The imaging path employs two achromatic doublet tube lenses (400–700 nm AR-coated, f=100mmm, Thorlabs) to relay the objective image onto the camera sensor. When combined with a 200 mm objective focal length, it reduces the image size by half, enabling an effective 0.5× demagnification of the sample onto the sensor.

A micromirror device (DMD) functions as a high-speed, digitally controlled spatial light modulator for patterned illumination of brain tissue. We use an achromatic Galilean beam expander (Thorlabs GBE02-A) to increase the size of the laser beam such that it hits a larger area of the DMD. To safely absorb laser energy not reflected by the DMD, we include a beam trap (Thorlabs BT610/M).

#### Excitation and emission bandpass filters

The excitation path includes the ET480/30x filter (center wavelength 480 nm, 30 nm bandwidth) to isolate blue light for fluorophore excitation. The emission path uses the ET525/36m filter (center wavelength 525 nm, 36 nm bandwidth) to selectively transmit the green fluorescence emission while blocking excitation light and background.

#### Objective Lenses

For our widefield voltage imaging system, the 20× Mitutoyo Plan Apochromat objective (MY20X-804) with a numerical aperture (NA) of 0.42 and a long working distance of 20 mm was used.

#### Patch cable

To connect the laser from the objective towards the implanted fiber a dual patch cable (BFYL2LS01, Thorlabs) was used. The patch cable is held by a holder in the optimal focal plane below the objective and connected using a mating sleeve to the implanted fiber.

### Second microscope setup

The 2nd microscope widefield setup is similar to the 1st microscope, except it used only LED illumination (blue: M470L5, yellow: SOLIS-565D, both from Thorlabs) instead of a laser and it includes an emission beam splitting component for dual color imaging. To allow simultaneous blue and yellow excitation, we used a multi-band dichroic (59022bs, Chroma). Emission wavelength splitting: The emission is effectively demagnified by combining a 1:1 Tube lens (TTL200, Thorlabs) and a 300mm focal length convex lens giving 0.6x demagnification. The emission is then split by a 570nm cut-off long-pass dichroic mirror (T570lpxr, Thorlabs). The emission paths are filtered respectively with 525/36mm and 630/75mm emission filters (both Chroma). With use of adjustable right-angle kinematic mirror mounts (KCB1C/M, Thorlabs), the green and red emission are projected such that they project onto separate nearby FOVs on the sCMOS. The emission paths are rejoined, while now shifted to each other, after with another 570nm cut-off long-pass dichroic mirror before reaching the sCMOS sensors. The illumination irradiance for Figure7 was 5-7 mW/mm^2^ for GEVI and 0.5-1mW/mm^2^ for mCherry.

### Animal experiments and surgical procedures

All procedures were approved by the appropriate institutional animal care and use committees and complied with relevant ethical guidelines. Experiments were performed on three to six months old female and male mice. The mice were maintained on a 12/12 h light/dark cycle and experiments were performed during the light cycle.

### Stereotaxic virus-injection surgery

Thirty minutes prior to the surgeries, analgesic (Buprenorphine, 0.05 mg/kg, s.c.) and anti-inflammatory medication (Carprofen, 5 mg/kg, s.c.) were administered. General anesthesia was induced in an induction chamber with 3 - 5 % isoflurane in 1 l/min oxygen. After the anesthesia induction the mice were head fixed in a stereotaxic frame, and the anesthesia maintained at 0.8–2% isoflurane via a nose cone. The body temperature was maintained at 37 °C using a feedback-controlled heating pad. To prevent corneal damage the eyes were covered with eye ointment (Duratears Z; Alcon, Netherlands) during the surgeries. After cleansing the scalp with alternating scrubs of chlorhexidine and 70% ethanol, the skin was excised. 2% lidocaine were applied to the exposed skull and then the periosteum was removed. The skull was aligned such that bregma and lambda were within 0.05 mm dorsoventrally and mediolaterally. Injection coordinates were marked for the CA1 (−2 AP, (−/+)1.8 ML, 1.12 DV from Bregma), mPFC ( 1.94 AP, 0.4 ML, 1.55 DV from Bregma) or striatum (−0.5 AP, −1.8 ML, 2.5 DV from Bregma) using a 25 G needle, and burr holes (∼0.25 mm) were drilled until the inner bone layer fractured. Glass micropipettes (30–40 µm tip diameter) were backfilled with mineral oil and mounted on a Nanoject III injector.

To allow Cre-dependent expression of the voltage indicator Ace-mNeon2 in PV-positive (PV+) neurons, PV-Cre-mice crossed with C57BL/6 mice were injected with the construct AAV9-CAG-DIO-Ace-mNeon2-Kv2.1PR (titer 2.79E+10). C57BL/6 mice were injected with the construct AAV9-CaMKII-pAce-Kv2.1PR, to allow the expression of the pAceR indicator in pyramidal neurons (titer 1.18E+11). The AAV constructs (CA1 pyramidal neurons: AAV9-CaMKII-pAce-Kv2.1PR co-injected with AAV5-hSyn-mCherry; CA1 parvalbumin neurons: AAV9-CAG-DIO-Ace-mNeon2-Kv2.1PR; ; striatal neurons: AAV9-CAG-DIO-Ace-mNeon2-Kv2.1PR co-injected with AAV.hSyn.Cre.WPRE.hGH) were diluted in sterile saline, and were loaded on a glass pipette. The pipette was slowly lowered to the final target depth. Injections were made at 1 nl/s. To minimize backflow, the pipette was left in place for 10 minutes post-injection before being slowly withdrawn. In some preparations we continued directly with the chronic fiber implantation after this step. The surgical area was irrigated with saline after the injection and then was dried thoroughly. At the end of the surgery the skin was sutured (Surgicryl 910) and sealed with a small amount of Vetbond if necessary. Following surgery, the isoflurane was discontinued and mice were monitored under 1 l/min oxygen until spontaneous movement resumed. Right after the mice were placed in a cage under a heating lamp and received sterile saline (1 ml, s.c.). Buprenorphine was administered after 8 hours of the first injection (0.05 mg/kg, s.c.). The body weight of the mice, the fur and the wound condition, as well as the general behavior were monitored daily for up to five days post-operatively. Any signs of distress were treated under veterinary guidance.

### Chronic Fiber Implantation surgery for Voltage Imaging

Two to three weeks after virus delivery (in some cases directly after the virus injection), mice underwent a second surgery to implant optic fibers, and a custom titanium head post. Buprenorphine (0.05 mg/kg, s.c.) and carprofen (5 mg/kg, s.c.) were administered 30 minutes before anesthesia induction. The anesthesia protocol, the surgical preparation, and the pre- and post-operative treatment were identical to the virus-injection surgery. After cleansing the scalp, the skin above the implantation area was removed, the skull was roughened and a support screw was placed anterior to the inferior cerebral vein, above the olfactory lobe. A burr hole was drilled at the previously marked location at the CA1 or striatum. Custom build implants containing an optic fiber (200um, 0.39 NA, RWD) and a twisted platinum-iridium bipolar electrode (101R-3T, A-M systems) were inserted at the CA1 or striatum and secured in place with C&B Metabond. Additional layers of Metabond and Tetric EvoFlow were applied to reinforce implant stability and ensure optical clarity. The titanium head post was placed posterior to the implant and anchored with Metabond, which was extended to cover the skull. Electrode wires were routed away from the fiber area, fixed in place, and encapsulated in dental composite. All implants remained stable and infection-free throughout imaging sessions.

### Imaging sessions

One week after surgery, animals were habituated to the head-fixed treadmill setup (Fig. 1a), where they were allowed to voluntarily run. Recordings were conducted for 1–3 weeks post-surgery. During head-fixed recordings, mice were head-fixed in the setup, and stimulation electrodes as well as LFP recording electrodes were connected. The optic fiber was coupled to a patch cable, and the holder was aligned with the focal plane of the objective. For data reflected in Figures 1-6 from the 1st microscope setup, a DMD pattern was projected onto the optic fiber to spatially pattern the laser light within the fiber. For Figure 7, based on the 2nd microscope setup, non-patterned LED was used for illumination. Video voltage imaging recordings from within the fiber were acquired simultaneously with LFP signals, pupil and body camera recordings, and treadmill velocity measurements. For freely moving recordings, the mice were briefly head-fixed to allow connection of the patch cable, stimulation electrodes, and LFP recording electrodes, after which they were placed in a recording cage.

### Deep brain stimulation

Electrical stimulation (Fig. 4-5, 7) was generated by a voltage generator (A-M Systems, Model3800) and delivered via an isolated pulse stimulator (A-M Systems, model 3820), which generated square-wave biphasic current pulses at either 40 Hz or 135 Hz. Peak stimulation amplitudes ranged from 10 to 100µA. Stimulation sequences were triggered by a MATLAB script through a National Instruments DAQ board, and the resulting pulses were recorded at a 30 kHz sampling rate using the Open Ephys platform.

### Data preprocessing

#### Synchronization of Optical and Electrical Recordings

To achieve precise temporal alignment between genetically encoded voltage indicator (GEVI) traces and local field potential (LFP) recordings, camera trigger pulses were digitized via an Open Ephys ADC channel. Imaging frames were registered to trigger timestamps by detecting rising edges in the ADC signal. LFP signals, acquired at 30 kHz, were downsampled to the imaging frame rate (∼500 Hz) by extracting samples corresponding to each camera trigger. For multi-trial paradigms, triggers were segmented into discrete trials using trial marker channels, with each trial processed independently.

### Fiber Photometry Preprocessing

Raw fluorescence traces were extracted from manually defined regions of interest (ROIs). Pixels corresponding to the fiber input face were included. To minimize potential optical edge artifacts and background noise, the ROIs were manually circumscribed only within the fiber core, specifically excluding the fiber-optic rim and cladding. To correct for photobleaching, traces were fitted with a double-exponential model applied to the pre-stimulation baseline. Corrected fluorescence Fcorr(t) was converted to ΔF/F using a static baseline method: ΔF/F = (F_corr - F0) / F0, where F0 was the mean fluorescence during the pre-stimulation period (typically 4 seconds for stimulation trials, or the entire trace for baseline trials). Narrow-band noise was removed using third-order Butterworth bandpass filters at 120–124 Hz and 130–132 Hz.

### LFP Preprocessing

LFP signals were recorded from contralateral hippocampal CA1 and, when available, from the ipsilateral hippocampal CA1. Signals were downsampled and aligned to camera triggers as described above. Outlier samples (∣z-score∣>10) were replaced with the median value.

### Artifact Removal

LFP artifacts were identified using a multi-stage pipeline combining median absolute deviation (MAD) thresholding (6 × MAD), rolling variance (4 × median variance over 0.5 s windows), and z-score thresholding (∣z∣>4). Artifact segments were first smoothed using morphological dilation with a 50 ms window to merge nearby artifacts, then extended by 100 ms on either side using padding. Segments shorter than 50 ms were discarded. In retained trials, artifactual samples were removed, producing discontinuous time series for subsequent analyses.

### Behavioral Classification

Running velocity was derived from rotary encoder signals (19.0 cm wheel diameter, 1024 counts per revolution) and smoothed using a 10-sample moving average. Behavioral states were classified using a stringent criterion: RUN periods required speeds >2.0 cm/s for a minimum bout persisting for at least 0.3 s, whereas REST periods required speeds <0.1 cm/s for ≥0.3 s. Intermediate velocities or bouts not meeting duration criteria were excluded from state-specific analyses to avoid contamination with transitional states.

### Coherence and Spectral Analysis

#### Coherence Estimation

Coherence between GEVI and LFP signals was estimated using two complementary approaches.

1. Welch-Based Estimation: Coherence was computed using MATLAB’s mscohere with Hanning-windowed segments. Baseline and overall stimulation recordings used 1.0 s segments with 80% overlap, while transient periods used 0.5 s segments with 50% overlap. Power spectral densities (PSD) were calculated via pwelch with identical parameters. Frequency resolutions were 2–70 Hz for baseline and 2–150 Hz for stimulation analyses.
2. Multi-Taper and FieldTrip Estimation: Coherence was validated using FieldTrip’s multi-taper method (mtmfft)^39^ with a single Hanning taper for both baseline and stimulation, enabling high-resolution detection of narrow-band peaks. Continuous data were segmented into pseudo-trials (1.0 s epochs, 80% overlap) for trial-averaged estimates via ft_connectivityanalysis with 0.5 Hz steps. Time-resolved coherence was visualized with sliding windows (baseline: 5.0 s window, 0.25 s step; stimulation: 0.5 s window, 0.05 s step). PSD is reported in decibels (10xlog_10_) and coherence as magnitude (0–1, with 0=uncorrelated and 1= perfectly correlated).

### Spectrograms and Spectral Normalization

Spectrograms were computed with MATLAB’s spectrogram function using overlapping windows (baseline: 0.75 s, 90% overlap; stimulation: 0.6 s, 88% overlap). Frequency resolution was enhanced via NFFT multipliers (2× for baseline, 3× for stimulation) and smoothed using 2D convolution. Power values were expressed in decibels (10xlog_10_) and baseline-normalized as fractional change for averaged stimulation trials: (P - P_baseline) / (P + P_baseline), where P_baseline was the mean power during the pre-stimulation period.

### Deep Brain Stimulation (DBS) Analysis

Stimulation onset was detected from ADC channels recording pulse train triggers sent by the voltage generator Model3800. Trials were segmented into pre-stimulation (–4 to 0 s), transient stimulation (0–0.15 s), sustained stimulation (0.15–1.0 s), and post-stimulation (1.0–5.0 s) intervals. For transient periods, window sizes were reduced to ensure multiple overlapping segments, and coherence values were cross-checked to minimize electrical artifact contributions. Band-specific power (35–45 Hz for 40 Hz DBS; 130–140 Hz for 135 Hz DBS) was extracted and baseline-normalized. Statistical comparisons across periods and stimulation frequencies were performed using paired t-tests or Wilcoxon signed-rank tests, with significance thresholds adjusted for multiple comparisons.

### Phase-Aligned Wavelet Spectrogram and Phase-Amplitude Curves

To visualize theta-phase-dependent spectral dynamics, we computed phase-aligned wavelet spectrograms and phase-amplitude curves. Instantaneous theta phase (5–9 Hz) was extracted from the LFP using complex Morlet wavelet decomposition (5 cycles) [30]. Time-frequency representations of both LFP and GEVI signals were obtained via continuous Morlet wavelet transform (5 cycles) across 2–100 Hz (95 linearly spaced frequency bins). Continuous data were segmented into two-theta-cycle epochs anchored at consecutive phase troughs (−π). Spectral magnitudes were sorted into 36 phase bins (10° width, −π to π) per theta cycle and averaged across artifact-free epochs, separately for running and resting states (see Behavioral Classification). Per-cycle averages were concatenated for a two-cycle display. Phase-amplitude curves were constructed by computing phase-triggered averages of the theta waveform (5–9 Hz, 4th-order Butterworth, zero-phase) and low-gamma amplitude (mean wavelet magnitude, 30–60 Hz) across the same bins; grand means and 95% confidence intervals (mean ± 1.96 × SEM) were derived across epochs.

### Cross-frequency coupling (CFC)

Coupling strength was quantified using the modulation index (MI)^29^. For each running epoch, the theta phase (5–9 Hz) from the LFP and low-gamma amplitude (30–60 Hz) from the wavelet spectrogram were sorted into 18 phase bins (20° each). The mean amplitude per bin was normalized to a distribution, and MI was defined as the Kullback–Leibler divergence from uniformity, normalized by log(N_bins_). MI was computed from all running epochs concatenated across trials. Statistical significance was assessed via iterative amplitude-adjusted Fourier transform (IAAFT) surrogate testing^40^. One thousand surrogates were generated by replacing the amplitude time series within each epoch with an IAAFT surrogate (50 refinement iterations), preserving the original power spectrum and amplitude distribution while disrupting phase-amplitude relationships. A one-sided p value was computed as (k + 1)/(N + 1), where k is the number of surrogate MI values ≥ the observed MI.

## Acknowledgement

This study was supported by the ERC starter grant, 101116609 (E.L.) and ParkinsonFonds Stichting grant,1914 (E.L.). We thank Jianguang Ni for comments and the whole lab for support.

