## Supplemental Figures 1-2 for "Tracking multi-site somatic voltage dynamics via high-speed fiber photometry"

### Supplementary materials

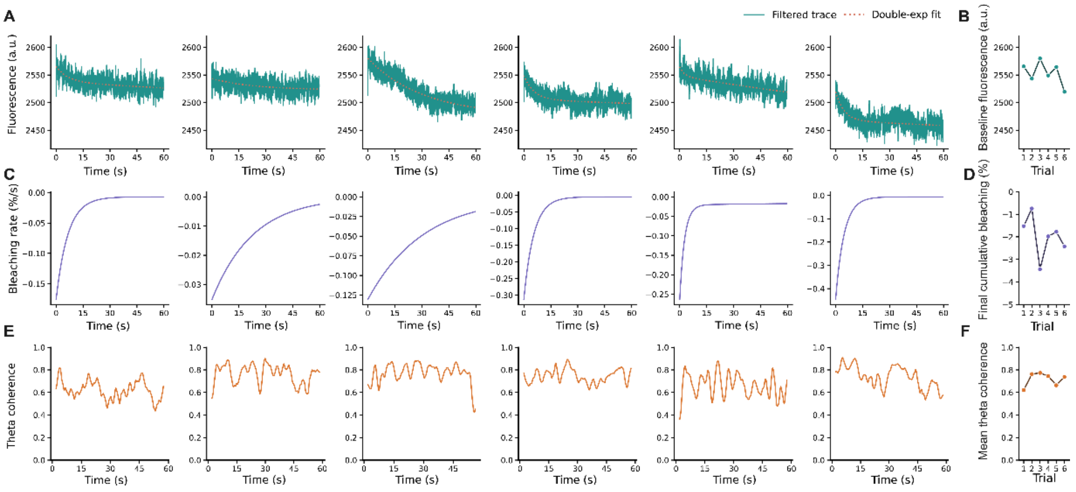


**Supplementary Figure 1. Photobleaching characterization** *(A) Raw fluorescence (pAce-KV2.1, green line) of 6 consecutive trials of 1min, separated by 5min. Dashed line represents the double exponential fit. (B) The mean fluorescence for the 6 trials. (C) The estimated bleaching rate (fluorescence change in percentage per second, %/s) for the 6 trials. (D)The mean bleaching rate for the 6 trials. (E) The estimated theta coherence between GEVI and LFP over time for the 6 trials. (F) The mean GEVI-LFP theta coherence for the 6 trials.*

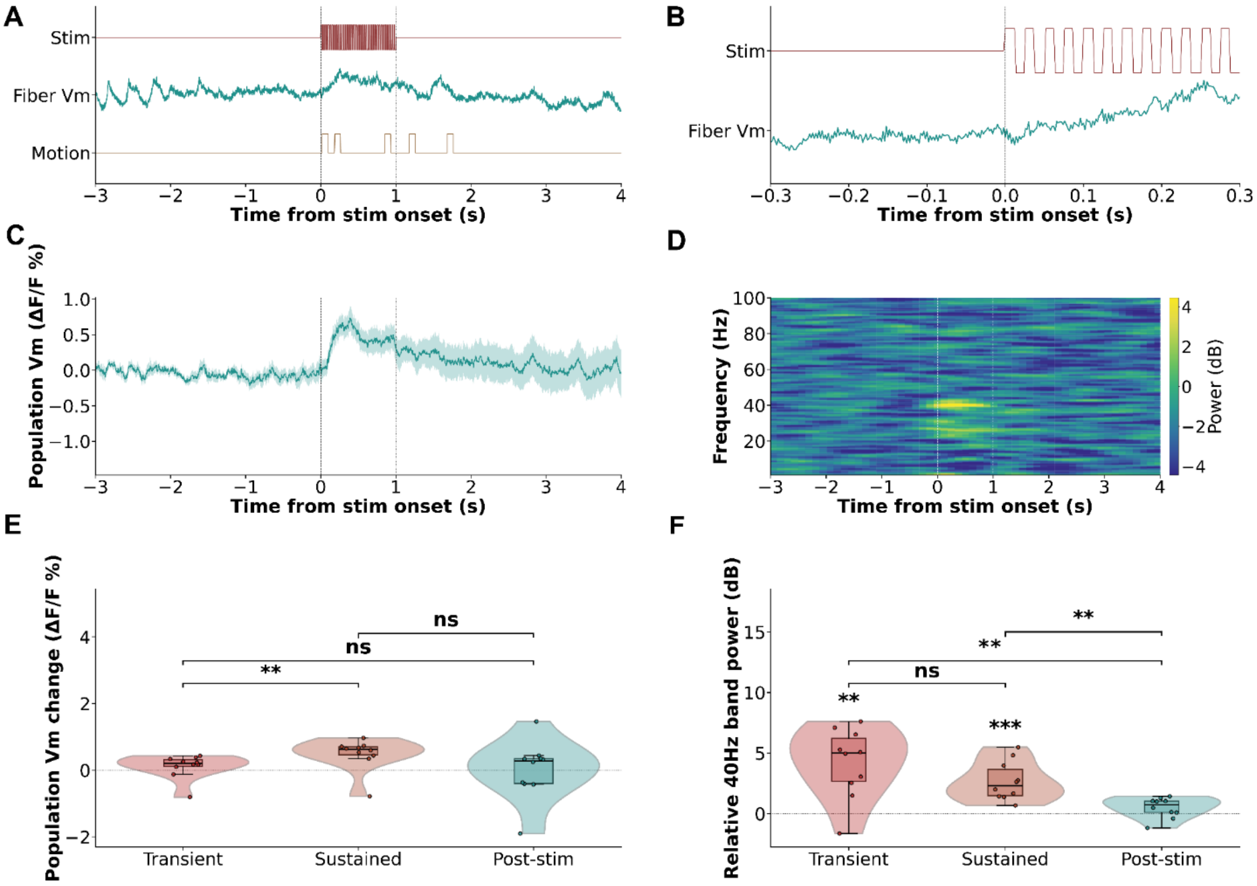


**Supplementary Figure 2. DBS modulated striatal voltage signals measured by GEVI fiber photometry.** *(A, B) Representative single trial recordings showing GEVI fluorescence (top, teal), and movement artifacts signals (bottom, orange) during 40Hz DBS (A) as well as a zoomed vies of the same trace (B, left). top bars indicate stimulation periods. (C) Average GEVI fluorescence traces (mean ± s.e.m; top) showing distribution across trials. (D) Time-frequency spectrogram during DBS computed using a 0.6s sliding window with 88% overlap and shown as relative power changes from baseline. (E, F) Violin plots showing population Vm changes across trials (left) as well as relative 40Hz band power (right).*
